# The genome of a sea spider corroborates a shared Hox cluster motif in arthropods with reduced posterior tagma

**DOI:** 10.1101/2024.11.20.624475

**Authors:** Nikolaos Papadopoulos, Siddharth S. Kulkarni, Christian Baranyi, Bastian Fromm, Emily V.W. Setton, Prashant P. Sharma, Andreas Wanninger, Georg Brenneis

## Abstract

**Background:** Chelicerate evolution is contentiously debated, with recent studies challenging traditional phylogenetic hypotheses and scenarios of major evolutionary events, like terrestrialization. Sea spiders (Pycnogonida) represent the uncontested marine sister group of all other chelicerates, featuring a – likely plesiomorphic – indirect development mode. Accordingly, pycnogonids hold the potential to provide crucial insight into early chelicerate genome evolution and ancestral principles of chelicerate body patterning. Leveraging this potential, however, has hitherto been hampered by the lack of high-quality genomic resources for pycnogonids.

**Results:** We employed long-read sequencing and proximity ligation data to assemble the first near chromosome-level sea spider genome for *Pycnogonum litorale*, complemented by developmental transcriptomes and a high-fidelity Iso-Seq dataset. The assembly has a size of 471Mb in 57 pseudochromosomes, a repeat content of 61.05%, 15,497 predicted protein-coding genes, and is highly complete (95.8% BUSCO Arthropoda score, 95.7% of conserved microRNA families present). We identified a single, intact Hox gene cluster that lacks *abdominal-A/Hox9*, suggesting the loss of this Hox gene.

**Conclusions:** Our high-quality genomic and transcriptomic resources establish *P. litorale* as a key research organism for modern studies on chelicerate genome evolution, development, and phylogeny. The presence of a single Hox cluster in the *P. litorale* genome further strengthens the inference that no whole-genome duplication occurred at the base of the chelicerate tree. The lack of *abdA* suggests that the combination of *abdA* loss and posterior tagmata reduction is a common theme in arthropod evolution, as it is shared with other, distantly related arthropod taxa with a vestigial opisthosoma/abdomen.

## Introduction

Chelicerata represents an extremely diverse arthropod lineage boasting more than 120,000 extant species (Brusca et al., 2023). They inhabit a wide range of habitats and have adopted highly divergent life strategies, as impressively evidenced by today’s terrestrial arachnid taxa (such as spiders, scorpions, harvestmen, mites and ticks). In contrast to most of their chelicerate kin, horseshoe crabs (Xiphosura) and sea spiders (Pycnogonida), plus selected mite taxa, are the only extant groups that inhabit the world’s oceans (Giribet & Edgecombe, 2019; Pepato et al., 2022).

Paramount to the chelicerate radiation has been the evolutionary plasticity of their body plan. One of its widely conserved hallmarks is the presence of two tagmata: the prosoma and the opisthosoma (Sharma & Gavish-Regev, 2024). The anterior prosoma comprises the ocular region and typically (but not always) six segments bearing the eponymous raptorial chelicera, followed by the pedipalp and four pairs of legs, all of which have been subject to considerable transformations in different lineages. By contrast, the posterior opisthosoma displays far greater evolutionary plasticity across chelicerates, not only in terms of segment number (up to 12 plus the asegmental terminus, the telson), but also with regard to the presence and function of diverse appendage derivatives (e.g., book gills, book lungs, or spinnerets) (Dunlop & Lamsdell, 2017; Sharma & Gavish-Regev, 2024).

The body organization of sea spiders is a unique variation on the chelicerate theme, characterized by several lineage-specific traits. The prosoma carries a prominent suctorial apparatus (=proboscis), a heavily modified first leg (=oviger) used for egg-carrying and grooming, and at least four pairs of true walking legs (Arnaud & Bamber, 1987). However, three distantly related pycnogonid taxa even possess five or six leg pairs (Ballesteros et al., 2021; Hedgpeth, 1947; Sabroux et al., 2023), which showcases a variability in the segmental composition of the prosoma that is uncharacteristic for the other chelicerate taxa. On the other hand, the opisthosoma of pycnogonids is dramatically reduced and represents only a small posterior protrusion (anal tubercle or “abdomen”) (Fig. 1A). Notably, it remains unclear to what extent vestigial opisthosomal segments contribute to the anal tubercle (e.g. Brenneis et al., 2023).

**Fig. 1:**
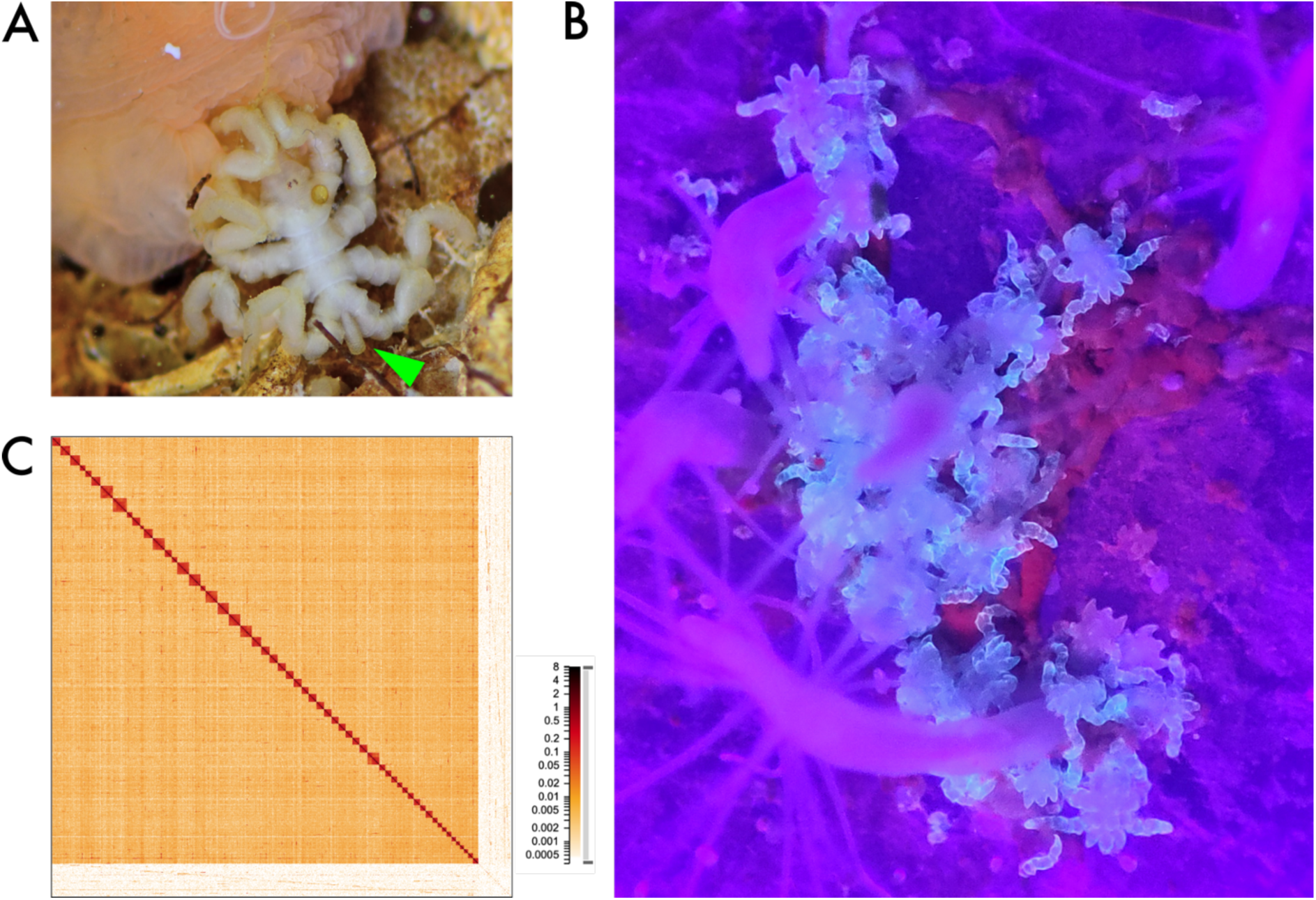
A) Adult *Pycnogonum litorale* specimen, feeding on the sea anemone *Metridium senile*. The green arrowhead points to the vestigial opisthosoma/anal tubercle. B) Post-embryonic instars V of *P. litorale* (fluorescing in light blue under UV light exposure), feeding on the hydrozoan *Clava multicornis*. C) Chromatin contact map, generated from Omni-C data at 1kb resolution, showing the 57 pseudochromosomes of the haploid genome (dark squares) and the unscaffolded contigs (bottom right region). Color denotes the number of contacts found in each region.

Notwithstanding their peculiar adult body organization, pycnogonids are the only extant chelicerates that display a pronounced indirect development with a segment-poor primary larva, a trait that may not only be plesiomorphic for chelicerates but also for arthropods in general (Liu et al., 2016; Waloszek & Maas, 2005; Wolfe, 2017). Accordingly, the study of sea spider development holds the potential to crucially inform debates on the evolutionary trajectories of chelicerate body patterning (Brenneis et al., 2017; Schwager et al., 2015). This is further underscored by their position in the chelicerate tree of life. After a long phylogenetic odyssey (Dunlop & Arango, 2005), pycnogonids are now robustly established as sister group of all other taxa (=Euchelicerata), rendering them one of the few stable anchors in the historically contentious and still controversial higher-order phylogeny of chelicerates (Ballesteros et al., 2022; Ballesteros & Sharma, 2019; Garwood & Dunlop, 2023; Lozano-Fernandez et al., 2019; Regier et al., 2010; Sharma et al., 2014).

This stable pycnogonid-euchelicerate sister group relationship also highlights a key role of sea spiders in interpreting the evolution of chelicerate genomes. For instance, the recently proposed hypothesis that whole-genome duplication (WGD) events in arachnopulmonates (spiders, scorpions and kin) and in xiphosurans (Ontano et al., 2021; Schwager et al., 2017; Sharma, 2023b; Shingate et al., 2020) represent derived states within chelicerates (e.g. Sharma, 2023b) is based on the lack of extensive duplications in so-called apulmonate taxa (harvestmen, ticks, mites) (Brückner et al., 2022; De et al., 2023; Gainett et al., 2021; Grbić et al., 2011). However, with euchelicerate interrelationships still in flux, polarization of WGD events and the reconstruction of the ancestral chelicerate condition remain challenging without a high-quality genomic resource for pycnogonids that can be scrutinized for the extent of gene duplication signatures. To date, the availability of but a single scaffold-level draft genome for *Nymphon striatum* (Jeong et al., 2020) and a handful of developmental transcriptomes (Ballesteros et al., 2021) has prevented inclusion of pycnogonids in macroevolutionary comparative studies. This includes studies on the composition of the Hox gene cluster, one of the best-known and most extensively discussed syntenic motifs of metazoan genomes (e.g. Garcia-Fernandez, 2005). In euchelicerates, Hox gene cluster duplications served as the first strong indicators for WGD events (e.g. Schwager et al., 2017). Functionally, Hox genes play a crucial role in the specification of segment identity along the anterior-posterior body axis of arthropods. This renders them crucial targets in the study of the genetic underpinnings of some of the unique features of the pycnogonid body plan.

The sea spider species *Pycnogonum litorale* (Strøm, 1762) from the north Atlantic is an emerging laboratory organism that allows to bridge this knowledge gap. It can be kept in laboratory cultures (Fig. 1A, B), is long-lived and displays year-round reproduction with a few thousand eggs per mating (Tomaschko et al., 1997; Wilhelm et al., 1997). In addition to these favorable characteristics, a solid body of morphological data on ontogenesis and adult morphology are available for this species (e.g. Alexeeva & Tamberg, 2021; Machner & Scholtz, 2010; Ungerer & Scholtz, 2009; Vilpoux & Waloszek, 2003). As a result, it has contributed to a number of developmental genetic investigations, both in standalone works as well as comparative studies (Brenneis et al., 2018, 2023; Klementz et al., 2024; Scholtz & Brenneis, 2016). To complement this morphological foundation, we present here the first genome of *P. litorale* together with transcriptomes of various developmental stages of its life cycle.

## Material and Methods

### Animal collection and husbandry

The *P. litorale* specimens used for genome and RNA sequencing originated from two different geographic localities: The first draft genome assembly is based on a single adult female from our *P. litorale* culture at the University of Vienna, Austria (specimen H1). This laboratory population derives from animals collected during low tide in the rocky intertidal of Helgoland in the North Sea (see (Brenneis et al., 2023) for details on subsequent husbandry). All developmental stages (embryonic stages, postembryonic instars I-VI, first juvenile instar and subadult) used for RNA-seq were reared and collected from the same in-house lab population.

For Omni-C and PacBio sequencing, *P. litorale* specimens were collected in the Gulf of Maine off the US East coast under rocks in tide pools at low tide by Tim Sheehan (44°54’03.5”N 67°07’17.8”W, Pembroke, Maine, United States) and transported to Madison, Wisconsin. This wild population constitutes the source of previously published transcriptomic data for *P. litorale* (NCBI <u>SRR7879353</u>). One male specimen (specimen M1) was used for PacBio sequencing and the second (specimen M2) for Omni-C sequencing.

We confirmed species identity of the two populations via cytochrome *c* oxidase subunit I (COX1) barcoding with widely used standard primers (Folmer et al., 1994). Obtained sequences were aligned and compared with corresponding barcodes of our in-house lab population and of additional *P. litorale* sequences deposited in the NCBI repository. Sequence identity ranged from 98.6 to 100% across all samples included in the alignment (Suppl. Fig. 1, Suppl. Table 2). Between the Gulf of Maine specimens and the lab population from Helgoland, sequence identity was even higher (99.5-100%), confirming the widely accepted trans-Atlantic distribution of *P. litorale* previously inferred from morphological species determination (e.g. Bamber, 2010).

A detailed table of all BioSamples involved in this study can be found in Suppl. Table 1.

### High molecular weight (HMW) DNA extraction and sequencing (ONT)

An adult female (specimen H1) was separated from the other animals and starved for a week in a small cage inserted in our seawater system without access to prey. Prior to DNA extraction, the animal was repeatedly rinsed in filtered artificial seawater (32‰, Red Sea, Red Sea Fish Pharm LTD) to minimize contamination by non-target DNA from externally attaching microorganisms. A Qiagen MagAttract HMW DNA kit (Cat. No. 67563) was used for DNA extraction following the manufacturer’s protocol, with the entire animal serving as input tissue. For this purpose, the female was placed in a Petri dish, cooled down on ice, quickly cut into several pieces using sterile microsurgical scissors and immediately transferred into the lysis buffer. After brief manual homogenization with a sterile pestle, the tissue pieces were incubated overnight at 56°C under constant shaking at 900 rpm. The HMW DNA was eluted in 100 µL AE buffer (10mM Tris-HCl, 0.5mM EDTA, pH 9.0).

The HMW DNA was subsequently transferred to the Vienna BioCenter (VBC) sequencing facility, where library prep and sequencing took place. Sequencing was performed on a PromethION machine. Reads were basecalled with the GPU-version of Guppy v6.5.7 (minimap2 v2.24-r1122 (Li, 2021)), using the r10.4.1 basecall model. Quality control was performed with nanoplot (De Coster & Rademakers, 2023).

### HMW DNA extraction and sequencing (PacBio)

Specimen M1 was maintained at 8°C without food for 11 days to minimize gut content contamination. High molecular weight DNA was extracted using the Qiagen MagAttract HMW DNA kit, following manufacturer’s instructions. After cooling on ice, the entire male was used as input tissue, with sectioning of live tissue with a sterile razor blade and immediate transfer into lysis buffer. After brief manual homogenization with a sterile pestle, the tissue pieces were incubated for 5 hr at 56°C with manual perturbation every 45-60 min, until the buffer appeared translucent. HMW DNA was eluted in 100 µL AE buffer. The beads were incubated in another 100 µL AE buffer overnight and eluted the following day. Samples were submitted for sequencing at the UW-Madison BioTechnology Center core facility on a PacBio Sequel II machine, using standard manufacturer’s protocols for the Sequel II Sequencing Kit 2.0. The library was sequenced on 1 SMRT Cell (8 M) in CCS mode for 30 h. Analysis was performed with SMRT Link v10.1 software, requiring a minimum of three passes for CCS generation.

### Omni-C sequencing

Specimen M2 was maintained at 8°C without food for 11 days to minimize gut content contamination and flash frozen with liquid nitrogen. The whole specimen was immediately transferred to dry ice and submitted for library preparation by Cantata Bio (Scotts Valley, CA, USA) with the Dovetail Omni-C library kit. Tissue from nearly the whole specimen was used to generate the Omni-C library. Chromatin was fixed in place with formaldehyde in the nucleus. Fixed chromatin was digested with DNase I and then extracted, chromatin ends were repaired and ligated to a biotinylated bridge adapter followed by proximity ligation of adapter containing ends. After proximity ligation, crosslinks were reversed and the DNA purified. Purified DNA was treated to remove biotin that was not internal to ligated fragments. Sequencing libraries were generated using NEBNext Ultra enzymes and Illumina-compatible adapters. Biotin-containing fragments were isolated using streptavidin beads before PCR enrichment of each library. The library was sequenced on an Illumina HiSeqX platform to produce ∼30x sequence coverage.

### RNA extraction and short-read sequencing

Embryonic stages, postembryonic instars I-VI, the first juvenile instar and subadults of *P. litorale* were separately transferred into Eppendorf tubes containing RNAlater (Sigma-Aldrich, Cat. No.: R0901) at ambient temperature. After the specimens had settled to the bottom, the tubes were left over night at 4°C and subsequently stored at −20°C until further processing. Extraction of total RNA was performed with a Qiagen RNeasy Plus Mini Kit (Cat. No.: 74134) according to the manufacturer’s protocol, eluted in 30 µL RNase-free water and stored at −70°C until sequencing. The RNA was transferred to the VBC sequencing facility, where library prep and sequencing took place. The libraries were sequenced on half of a NovaSeq S4 lane (300 cycles). The sequencing data was demultiplexed with RTA v3.4.4 and demultiplexed by bcl2fastq2 v2.20.0 with default parameters by the VBC sequencing facility.

Zygotes, early cleavage, and more advanced embryonic stages from our in-house colony were also sequenced independently in Madison, Wisconsin. The samples were preserved and transported in tubes containing RNAlater, and extracted using TRIzol reagent (Thermofisher, Cat. No.: 15596026) following manufacturer’s instructions, with elution into THE RNA Storage Solution (Thermofisher, Cat. No. AM7001). Library preparation with Illumina TruSeq library kits was performed at the UW-Madison BioTechnology Center. Paired end 2 × 150 bp sequencing was performed on an Illumina NovaSeq platform.

For an overview of samples and sequencing depth, please refer to Suppl. Table 1.

### Full length mRNA sequencing

Prior to enzymatic shearing, small volumes of the same RNA extractions used for short-read RNA sequencing were mixed to obtain an RNA pool covering all phases of development (Suppl. Table 3). The pooled sample was subsequently transferred to the VBC sequencing facility for long-read mRNA sequencing (Iso-seq) on a PacBio platform.

The data were processed according to the instructions outlined in the <u>publicly available</u> <u>documentation from PacificBiosciences (</u>https://isoseq.how/). Briefly, subreads were processed to circularized consensus sequences with ccs v6.4.0; the CCS were demultiplexed and primers were removed with lima v2.9.0; the resulting BAM files were refined by trimming poly(A) tails and removing concatemers with the refine command of the isoseq package (v4.0.0); the refined transcripts were clustered with the cluster2 command of the same package; the bam2fastq utility (v3.1.1) was used to convert the BAM files to FASTQ format; finally, the clustered transcripts were mapped to the genome with pbmm2 (v1.13.1) and the result was used to collapse the refined CCS reads with the collapse command of the Iso-seq package and obtain a GFF file. More details can be found in the GitHub repository for this study (https://github.com/galicae/plit-genome under 05-transcriptomes/).

### Genome assembly, scaHolding, and contaminant filtering

To estimate genome size and heterozygosity, we quantified frequency spectra for k-mers of size k=21 with jellyfish v2.3.0 (Marcais & Kingsford, 2012) and analyzed them on the GenomeScope (Vurture et al., 2017) and GenomeScope2 (Ranallo-Benavidez et al., 2020) webservers. This approach is intended for very highly accurate short reads, therefore the analysis of the long-read data should be viewed with caution, but with high enough coverage the k-mer spectra of error-prone reads should be approximating the true distributions.

The PacBio data had a k-mer coverage of 8-17x according to GenomeScope (Suppl. Fig. 2A, C), below the theoretical 30x threshold that was often suggested in the short-read era (Sims, 2014). Nevertheless, we tried an assembly with Flye v2.9.2 (Kolmogorov et al., 2019), using default parameters. The ONT reads had a 31x k-mer coverage. We tested Flye v2.9.2, shasta v0.11.1 (Shafin et al., 2020), and Verkko v2.0 (Rautiainen et al., 2023), all with default parameters.

To scaffold, we followed the “HiC map7” pipeline from Schultz, 2023, adapting it to our computing environment. We mapped the omni-C data onto the assembly using chromap v0.2.6-r490 (Zhang et al., 2021). The resulting SAM file was sorted and indexed with samtools v1.16.1 (using htslib 1.16) (Danecek et al., 2021), before being passed on to yahs v1.2a.2 (Zhou et al., 2023). The result was edited manually using juicebox v2.15 (Dudchenko et al., 2018; Durand et al., 2016).

All the scripts for genome assembly, quality control, and evaluation can be found in the GitHub repository (https://github.com/galicae/plit-genome under 01-assembly/).

We used MMseqs2 v6f45232 (Steinegger & Söding, 2017) to download the UniRef90 database (Suzek et al., 2015), index the draft genome for alignment, and align the draft genome against UniRef90, adding the taxonomic information of each hit. We summarized the results by aggregating the hits within each scaffold according to their taxonomic assignment. We then removed scaffolds where less than 90% of hits were of metazoan origin. This filtering marked and removed pseudochromosomes 52 and 53 as contaminants. However, we initially proceeded without renaming the following scaffolds, to be able to follow potential issues back to their original source. After the annotation procedure was finished, we used the Linux bash utility sed to rename pseudochromosomes 54-59 to 52-57, respectively, in the genome sequence (FASTA) and annotation (GFF) file. Bash scripts and Jupyter notebooks for the individual steps can be found in the GitHub repository (https://github.com/galicae/plit-genome under 04-contam/).

### Genome annotation

We used RepeatModeler v2.0.5 (Flynn et al., 2020) to build a repeat family database for *P. litorale*, running it with the -LTRStruct argument to try and characterize long terminal repeats. The families predicted by RepeatModeler were then used by RepeatMasker v4.1.6 to softmask the draft genome.

We used BRAKER v3.0.6 (Gabriel et al., 2024) with the short-read transcriptomic data and the Arthropoda OrthoDb v11 (Kuznetsov et al., 2023) protein sequences to predict protein-coding gene models on the softmasked draft genome. We used the default Augustus configuration directory, copied from a local clone of the Augustus repository (commit d0b1b6c, Nov. 27, 2023). We used the containerized version of BRAKER, run with Singularity v3.8.6. To speed up calculations, the transcripts were mapped to the draft assembly with STAR v2.7.11b (Dobin et al., 2013) before running BRAKER.

We merged the GFF file produced from BRAKER with the one produced from Iso-seq collapse (see Methods) with the intersect program of the bedtools suite (v2.30.0) (Quinlan & Hall, 2010) and used custom Python code to analyze the overlap (https://github.com/galicae/plit-genome under 06-annot/annot1-isoseq_confirm.ipynb). We tested how many Iso-seq clusters (interpreted as genes) were overlapping with each BRAKER gene model. We considered two cases of significant overlap for gene models and clusters that occupy similar loci on the same strand: first, if one gene model only contained transcripts from the same cluster, we asked that at least 40% of the transcripts either contained the full gene model or were fully contained within it. If that was not the case, we demanded that the average overlap between transcripts of the cluster and the gene model exceeded 50%. These cutoffs ensured that we did not merge a gene model with a cluster in cases of partial overlap (e.g. the 5-prime end of the gene model overlaps with the 3-prime end of the cluster). We generated a merged GFF file that retained the unique BRAKER gene models and Iso-seq clusters and reconciled the overlapping regions. Gene models deriving from the Iso-seq clusters were given the cluster name (“PB.number>”); gene models deriving from BRAKER kept their BRAKER name (“g <NUMBER>”).

We then used bedtools intersect with the -v flag to exclude from the RNA-seq data everything mapping to loci covered by the merged GFF. The reduced RNA-seq data were then supplied to BRAKER and a second round of gene model prediction was run with the same parameters as the first. Using bash command line tools and bedtools intersect, we extracted the exons from the round 2 GFF file and cross-referenced them with the merged GFF file; we used this information to extract novel gene models from the round 2 GFF file. These were then deposited in a new GFF file with the agat_convert_sp_gxf2gxf.pl script (AGAT suite) and appended to the merged GFF. To avoid confusion, novel gene models kept their BRAKER-style IDs but with a prefix (“r2_g <NUMBER>”).

In a third step, we performed de novo assemblies for the deeply sequenced RNA-seq datasets (see Suppl. Table 1) with Trinity v2.15.1 (Grabherr et al., 2011), with the --trimmomatic and --no_salmon flags. We extracted complete ORFs from the de novo transcriptomes with TransDecoder v5.7.1 (Haas, 2023), mapped them against the draft genome with minimap2 v2.28, and exported the mapping results in GFF format with the agat_convert_minimap2_bam2gff.pl script from the AGAT suite (v1.4.0) (Dainat, 2024). We then used bedtools intersect with the -v flag to extract map events that did not overlap with the already annotated gene models (merged+round2 GFF). We then used bedtools to find the pairwise intersections of the resulting GFFs with the instar III-based GFF. The resulting overlap table was analyzed with custom Python code (see GitHub repository, https://github.com/galicae/plit-genome under 06-annotation/) and the overlaps were reconciliated into gene models, which were given ascending unique IDs similar to Trinity transcript IDs (“DN <NUMBER>”) but with a prefix to denote their origin (“at_DN <NUMBER>”, for “assembled transcriptome”). The result was written out in GFF format and concatenated to the merged+round2 GFF file. Finally, the merged GFF was sorted and formatted with genometools gff3 v1.6.2 (Gremme et al., 2013), producing the final protein-coding gene annotation. The entire process is described in-depth in the GitHub repository (https://github.com/galicae/plit-genome under 06-annotation/). Finally, we annotated tRNA genes with tRNAscan-SE v2.0.12 (Chan et al., 2021).

### Orthology assignment

We used the possible ORFs and agat_sp_extract_sequences.pl script from the AGAT suite (v1.4.0) to extract exons, using the draft genome and the final GFF file as input. The result was processed with TransDecoder v5.7.1 to produce a list of select the most probable ones. The TransDecoder-predicted peptide file, containing multiple entries per gene, was submitted to the EggNOG-mapper server (v2.1.12 Cantalapiedra et al., 2021) and mapped against version 5 of the EggNOG database (Huerta-Cepas et al., 2019) using default parameters. The results were downloaded and analyzed further with a Python notebook (https://github.com/galicae/plit-genome under 07-analysis/emapper_output.ipynb). To improve usability, annotations of different transcripts and isoforms were collapsed for each gene ID by keeping the entry with the highest bit score, leading to a slimmer look-up table.

### MicroRNA survey

For the prediction of the conserved microRNA complement and in the absence of small RNA sequencing, the genome was subjected to MirMachine (Umu et al., 2023) analysis, using protostome models and “Chelicerata” as search node based on MirGeneDB 3.0 annotations (Clarke et al., 2024). High and low confidence predictions were compared to the expected microRNA complement.

### Hox cluster annotation

To identify *P. litorale* Hox genes, publicly available Hox gene sequences from the sea spiders *Nymphon gracile* and *Endeis spinosa* were downloaded from NCBI (Sayers et al., 2022). In addition, the Hox gene sequences of the harvestman *Phalangium opilio*, an apulmonate representative with a well-annotated, unduplicated Hox gene complement, were used as queries in BLAST searches (tblastn) against our different *P. litorale* transcriptomes. The best *P. litorale* results (complete CDS) were used as queries in BLAST searches (blastx) against the NCBI database to confirm the assignment of Hox gene identity. The putative *P. litorale* Hox gene sequences were then used to scan the genome with MMSeqs2 using default parameters (https://github.com/galicae/plit-genome under 07-analysis/genes.sh).

For the posterior Hox gene *abdominal-A* (*abdA/Hox9*) we performed an extended, more targeted search against the draft genome (MMseqs2 with default parameters), using various spider *abdA* sequences as queries. More details can be found on the GitHub repository (https://github.com/galicae/plit-genome under 07-analysis/hoxfinder.ipynb).

### Code availability

All relevant code for data preprocessing, genome and transcriptome assembly, subsequent analysis, and figure generation is available at GitHub under https://github.com/galicae/plit-genome. An archived version of the code can be found on Zenodo (https://doi.org/10.5281/zenodo.14188290). The computational results of this work have been achieved using the Life Science Compute Cluster (LiSC) of the University of Vienna. At the time of writing, LiSC is operating Oracle Linux Server v9.4 and uses SLURM (Yoo et al., 2003) for the management of computational tasks. The code provided here is optimised for this environment but is easily adapted to any UNIX-based system.

The computing environments for certain tools were modularized as conda environments. A list of configuration files for the most recent environment per tool can be found in the repository as well.

### Data overview and availability

All sequencing data was deposited on ENA with the appropriate metadata (Study Accession PRJEB80537). An overview table can be found in Suppl. Table 1.

## Results

### Genome sequencing and assembly

Specimen H1 was sequenced to 30.5M reads (56.4Gb) with the ONT PromethION platform (N50: 5.3kb, average read length: 1.85kb). The M1 specimen was sequenced with the PacBio HiFi platform to 1.7M reads (10.1Gb). K-mer spectrum analysis on the GenomeScope (Vurture et al., 2017) and GenomeScope2 (Ranallo-Benavidez et al., 2020) webservers predicted medium levels of heterozygosity (approx. 0.68%). The predicted genome size was between 190-530Mb (Suppl. Fig. 2), a range below the only other currently available pycnogonid draft genome of *Nymphon striatum* at approx. 732Mb (Jeong et al., 2020). The low k-mer coverage of the PacBio reads (8-17x, see Suppl. Fig. 2A, C) discouraged us from using them for de novo genome assembly. Specimen M2 was sequenced to 79.4M reads (24Gb) using the Omni-C protocol and the Illumina platform. For an overview of sequencing data, see Suppl. Table 1.

To complement the genomic data, we generated 1.24B Illumina short transcriptomic reads (186.7Gbp) from 15 closely sampled developmental time points, spanning from the zygote to the sub-adult stage (Suppl. Table 1). We also generated 1.5M full length mRNA reads (3.43Gb) using the PacBio Iso-seq technology.

We tried a variety of de novo genome assemblers using both the ONT and HiFi data. A comprehensive table of alignment results can be found in the GitHub repository (https://github.com/galicae/plit-genome under 02-scaffold). The most promising assembly in terms of contiguity and completeness was produced with the ONT data using Flye. After scaffolding, manual curation of the Omni-C map, merging of smaller scaffolds by their Omni-C score, and scaffold decontamination (see Methods), the draft genome had a total size of 471.6Mb, with almost 93% of the sequences contained in 57 pseudochromosomes (in silico assembled pseudomolecules not verified experimentally) (Fig. 1C). The draft assembly reached a BUSCO (Manni et al., 2021) completeness of 95.8% (BUSCO Arthropoda) and an N50 of approx. 8Mb. The completeness and contiguity progress from initial assembly, through scaffolding, manual curation and decontamination is documented in Suppl. Table 4. The majority of genomic and transcriptomic raw reads mapped back to the draft assembly (ONT: 76.23%, PacBio: 84.38%, short RNA average: 87.75%, Iso-seq: 99.67%), indicating that the assembly captures the genetic variability of the laboratory culture and the wildtype specimen equally well.

### Genome annotation

#### Repeat annotation and analysis

Using RepeatModeler and RepeatMasker, a total of 61.05% of the assembly was identified as repetitive elements, with 11.65% being classified as retroelements (mostly LINEs, at 9.75%), 6.14% classified as DNA transposons, and 43.26% remaining unclassified (Fig. 2). As this level of repeat content is unusual for the modest genome size (see Fig. 3F), it prompted a deeper analysis. We first sought to confirm that the high amount of repetitive sequences was not an artifact of the genome assembly. To this end, we mapped the raw ONT and PacBio reads onto the repeat families predicted by RepeatModeler. We found close to 97 million matches on 18.5 million ONT reads (total: 30.5 million, for a 60.41% mapping rate), indicating that each read contained on average more than 5 repeats. The results were similar for the PacBio data, with 11 million matches on 1.37 million HiFi reads (total: 1.72 million, for an 80.1% mapping rate), indicating that each read contained on average 8 repeats. The high abundance of repeated sequences on the raw reads of two different sequencing technologies and specimens originating from different natural populations leads us to conclude that the annotated repeat content is likely not a prediction artifact. The repeat analysis results are available online in the GitHub repository (https://github.com/galicae/plit-genome under 04- contam/).

**Fig. 2:**
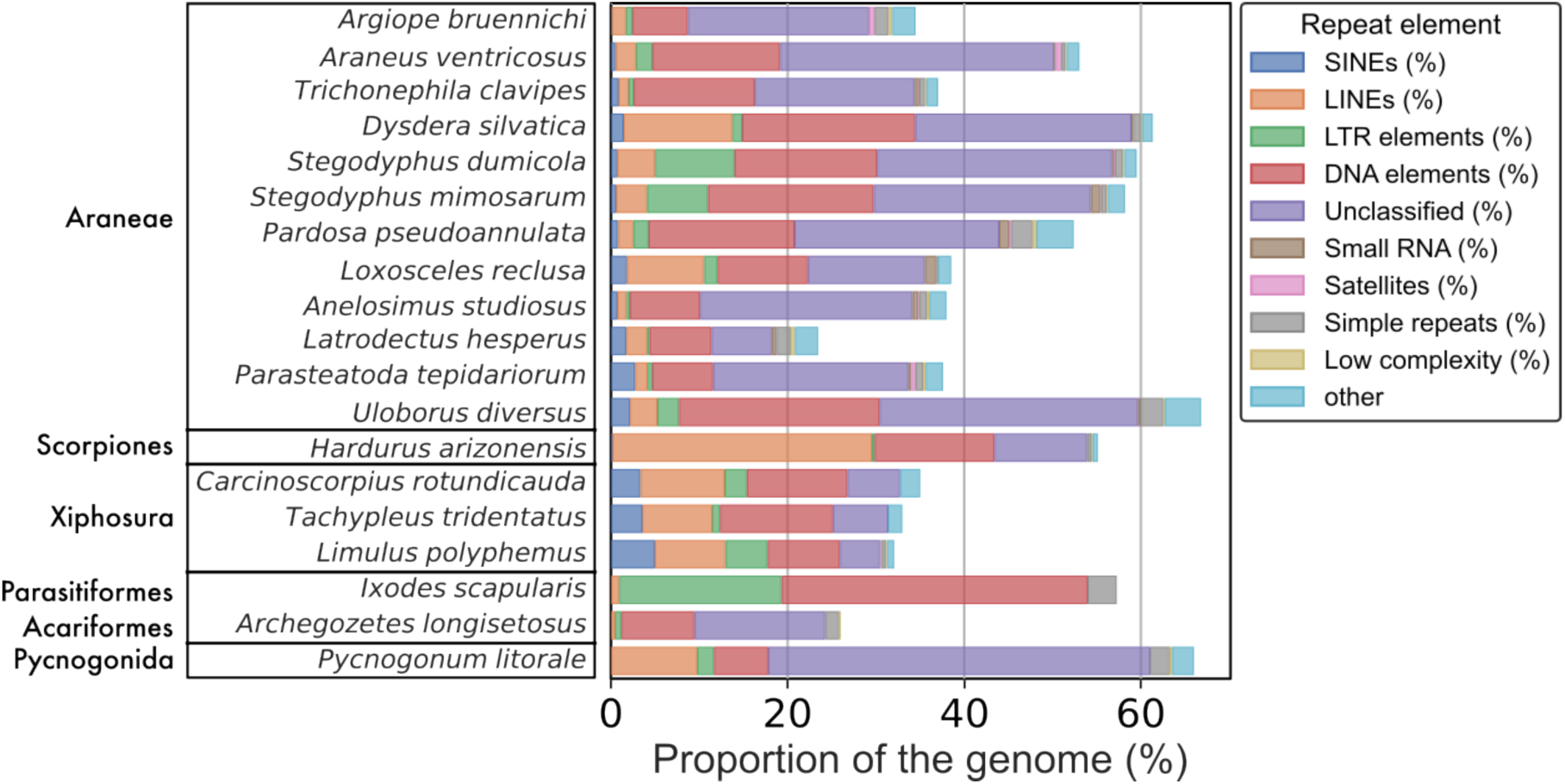
Breakdown of repeat content for various chelicerate genomes, including *Pycnogonum litorale*. Araneae as reported by Sheffer et al., 2021; *H. arizonensis* as reported in Bryant et al., 2024; Xiphosura as reported in Nong et al., 2021; *I. scapularis* as reported in Nuss et al., 2023; *A. longisetosus* as reported in Brückner et al., 2022. For underlying data, see Suppl. Table 9.

**Fig. 3:**
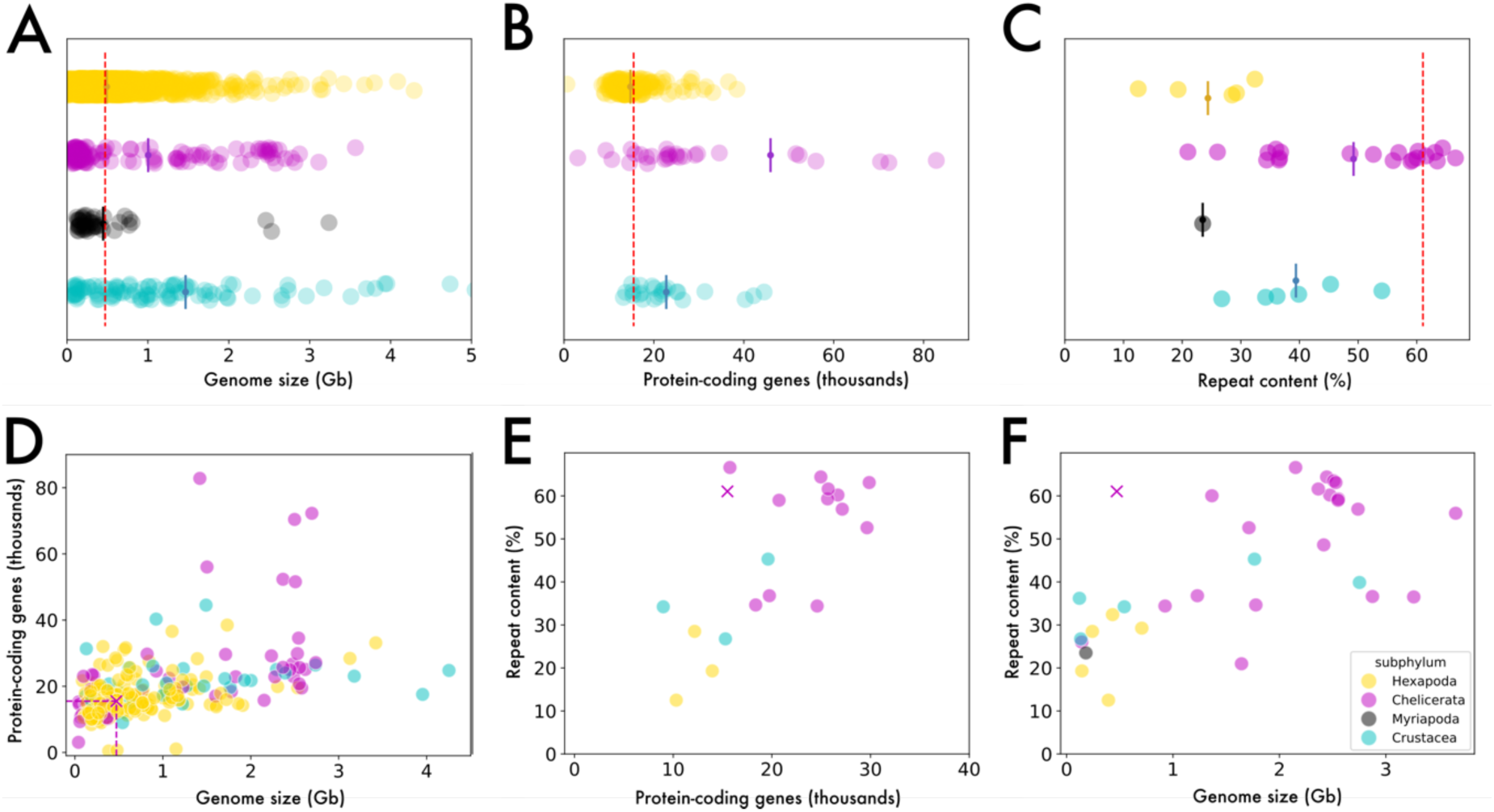
The *Pycnogonum litorale* genome in the broader arthropod genomic context. Each point represents one genome, with Hexapoda shown in yellow, chelicerates in magenta, myriapods in gray, and crustaceans in cyan. A)-C) Strip plots of the distributions of A) genome size, B) predicted protein-coding genes, and C) repeat content. The mean of each distribution is indicated by a bisected circle. The values for the *P. litorale* genome are indicated by a dashed red line. D)-F) Scatter plots showing the relationship of D) protein-coding gene number to genome size, E) repeat content to protein-coding genes, and F) repeat content to genome size. The values of the *P. litorale* genome are indicated by a cross. Data from NCBI (Suppl. File 1).

Next we investigated the unclassified repeats, as they made up the majority of the repeat content of *P. litorale.* Manual inspection of some unclassified repeat sequences retrieved low-confidence database hits (nucleotide BLAST against NCBI nr) from sea anemones, including *Metridium senile*, which is the prey of juveniles, subadults and adults of *P. litorale* in our laboratory culture. To exclude contamination as a possible explanation we identified 10,170 raw ONT reads (0.03% of total reads) that mapped with high confidence (Q>30) to both the *P. litorale* and *M. senile* genomes. Of these, more than half (4,593) are highly repetitive (>50% of read length) and another 14% (1,446) are mildly repetitive (>25% of read length). Furthermore, many of the affected *P. litorale* contigs are themselves highly repetitive, with one contig in particular containing 2,197 of the suspicious reads. Accordingly, the putative contaminants from *M. senile* are a vanishingly small number of highly repetitive reads on highly repetitive contigs (see also relevant notebooks in the GitHub repository under 04-contam/).

#### Gene prediction

Using the short-read transcriptome data for protein-coding gene prediction with BRAKER (Gabriel et al., 2024) yielded 11,451 gene models. However, suboptimal BUSCO scores (lower completeness, higher duplication rate) of the gene models (“protein” mode) compared to the entire genome (“genome” mode) prompted us to investigate the predictions. We cross-referenced this with the loci of the 8,904 clusters produced by the Iso-seq processing pipeline (isoform collapsing, see Material and Methods), and were able to identify multiple cases where BRAKER models of distinct transcript clusters were erroneously merged, suggesting that BRAKER did not detect the intergenic boundaries correctly. We also noticed a number of loci with RNA-seq peaks in characteristic exon-intron patterns without gene models, suggesting that, in these loci, the BRAKER annotation had been too conservative. We resolved these cases by treating Iso-seq transcript clusters as genes and ignoring BRAKER predictions that overlapped (same strand, any sequence overlap) with Iso-seq “genes”. This left 3,596 BRAKER gene models to complement the Iso-seq clusters, leading to a total of 12,500 gene models. We then filtered the short-read transcriptome data to exclude reads that mapped to loci with gene models and ran BRAKER again. This produced 2,223 gene models that did not overlap with the first round of annotation. We verified by manual inspection that many previously unidentified loci were now covered by gene models. Finally, we used the de novo assembled transcriptomes from deeply sequenced developmental time points to suggest another 754 putative gene loci, for a total of 15,497 gene models. The vast majority of genes (95.7%) are on the 57 pseudochromosomes, with only 670 gene models on smaller scaffolds. The completeness score for BUSCO Arthropoda remains stable (“genome” mode on the raw sequence of the draft genome: 95.8%, “protein” mode on the predicted proteome: 95.8%). A total of 10,017/15,497 (approx. 65%) of the gene models were assigned an orthogroup by EggNOG-mapper, meaning they mapped to known gene families. Finally, prediction of tRNA genes resulted in a total of 4189 genes, of which 965 were not overlapping with protein-coding genes.

#### MicroRNA content

Of the 47 conserved microRNA families predating the evolutionary origin of Chelicerata (1 Eumetazoa, 31 Bilateria, 11 Protostomia, 1 Ecdysozoa and 3 Arthropoda), we found 42 in the high-confidence predictions, indicating a very good performance of the Covariance models in MirMachine. Missing families included the bilaterian-specific Mir-242 and Mir-76 (with Mir-242 being a known loss in all Ecdysozoa), the protostome-specific Mir-1993 and Mir-67, and Iab-4. Manual investigation of the low-confidence candidates for these families identified predictions for Mir-76, Mir-1993, and Mir-67 with conserved seed sequences, likely representing true microRNAs. Hence, with 45 out of the 47 clearly expected families (95.7%), the genome assembly reaches a high completeness score for conserved microRNAs, serving as an additional indicator of its high quality. By contrast, out of the three microRNA families hitherto classified as being chelicerate-specific (Mir-3931, Mir-5305, and Mir-5735), none were recovered in the high-confidence predictions. However, a low-confidence prediction with a well-conserved seed sequence likely resembles a true positive for the Mir-3931 family. Of the predicted microRNA families, 11 had more than one copy, leading to a total of 70 predicted microRNA genes (Suppl. Table 10).

The intermediate and final GFF files are available on Zenodo (10.5281/zenodo.14185694); the final genome annotation is also deposited on the ENA server under accession ID XXX.

### The Hox gene cluster of *P. litorale*

Using the downloaded Hox gene queries we identified candidate sequences for nine of the putatively ten ancestral arthropod Hox genes in our *P. litorale* transcriptomes, namely *Labial* (*lab*/*Hox1*), *Proboscipedia* (*pb*/*Hox2*), *Hox3*, *Deformed* (*Dfd*/*Hox4*), *Sex combs reduced* (*Scr*/*Hox5*), *Fushi-tarazu* (*ftz*/*Hox6*), *Antennapedia* (*Antp*/*Hox7*), *Ultrabithorax* (*Ubx*/*Hox8*), and *Abdominal-B* (*AbdB*/*Hox10*). None of the queries yielded a candidate sequence for *abdominal-A* (*abdA*/*Hox9*). BLAST searches with the putative *P. litorale* Hox sequences against the NCBI database corroborated our preliminary Hox gene identification. Reverse translated search of these nine sequences against the draft genome recovered a single intact cluster spanning 1.07 Mb on pseudochromosome 56, with all hits overlapping gene models and no evidence for additional Hox clusters or single gene copies on any of the other pseudochromosomes. The nine Hox genes follow the typical ordering (ascending from *Hox1* to *Hox10*) and show identical orientation on the minus strand (Fig. 4A). By contrast, dedicated search of the genome with the putative but highly divergent partial *abdA* sequence of the sea spider *Nymphon gracile* (Manuel et al., 2006) did not result in any statistically significant hits (also see the “hoxfinder.ipynb” notebook on the GitHub repository under 07-analysis/). Given the lack of any other putative pycnogonid *abdA* sequences, we additionally searched the genome with various spider *abdA* sequences, which invariably matched either the locus of *Plit-Ubx* or of *Plit-Antp* (Suppl. Fig. 3; Suppl. Table 5), indicating that an *abdA* ortholog is truly lacking (or degenerated beyond recognition) in *P. litorale*. One minor irregularity in the cluster pertains to the presence of a putative gene between *Plit-Hox3* and *Plit-Dfd* (Fig. 4A). This gene model (r2_g3735) was produced in the second round of BRAKER annotation and a BLAST search against NCBI’s nr database finds excellent hits (alignment score ≥200) across very diverse taxa (fish, bivalves, sea anemones, sponges, bacteria; see Suppl. Table 6). However, all aforementioned BLAST hits are so-called “hypothetical” or “uncharacterized” proteins without any conserved domains, suggesting that the locus represented by r2_g3735 may be contamination, a jumping gene, or might represent a non-coding sequence.

**Fig 4:**
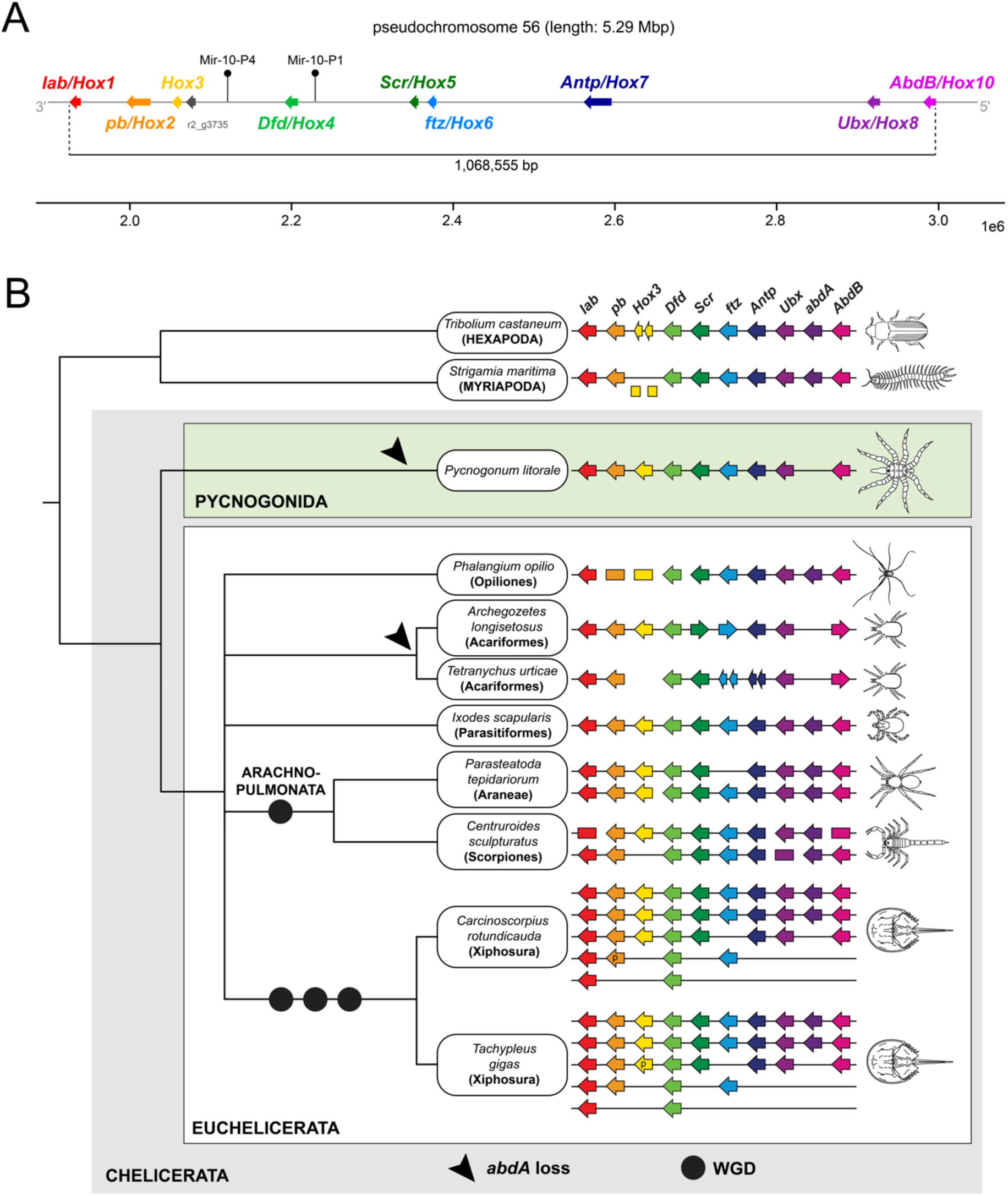
A) Schematic representation of the *Pycnogonum litorale* Hox gene cluster on pseudochromosome 56, arrows indicate the direction of transcription. Note the absence of *Plit-abdA* between *Plit-Ubx* and *Plit-AbdB*. The positions of the predicted microRNAs Mir-10-P4 and Mir-10- P1 are indicated by black pins. Note the absence of the positionally highly conserved Iab-4 between *Plit-Ubx* and *Plit-AbdB*. B). Hox gene clusters in chelicerate genomes and in selected outgroups (modified from Sharma, 2023a; data for *A.longisetosus* as reported in Brückner et al., 2022). Arrows represent the direction of transcriptional activity, where known. Circles represent whole genome duplication events (WGD), with at least two in the lineage leading to extant xiphosurans and another in the stem lineage of Arachnopulmonata. Arrowheads indicate the independent loss of *abdA* in Pycnogonida and Acariformes. p: pseudogene.

Two of the 70 predicted microRNA genes are located in the Hox cluster. They belong to the MIR-10 family, with MIR-10-P4 (previously named MIR-993) found between *Plit-Hox3* and *Plit-Dfd*, whereas the other paralogue MIR-10-P1 is located between *Plit-Dfd* and *Plit-Scr* (Fig. 4A).

## Discussion

### The *P. litorale* genome in an arthropod context

Genomic resources for Chelicerata are quite asymmetrically distributed, with gaps in high-quality datasets for several understudied orders (Garb et al., 2018). Prior to our study, a single pycnogonid draft genome of *Nymphon striatum* had been publicly available (Jeong et al., 2020). The small body size of this species led these authors to pool multiple (40) individuals, increasing the effective heterozygosity. This affected the quality of the final assembly that after scaffolding with mate-pair reads had an N50 of 701.8kb only, despite starting from PacBio HiFi reads with an N50 of 19.4kb. By comparison, our use of a single female of *P. litorale* as input material yielded a scaffold N50 an order of magnitude higher (7.97Mb), despite starting with ONT reads with an N50 four times lower (5.3kb). Even though both genomes have comparable BUSCO completeness scores (96-97% BUSCO Arthropoda), the *P. litorale* genome thus exhibits a significantly higher contiguity compared to the *N. striatum* assembly, reaching near chromosome-level.

The number of 57 pseudochromosomes in the *P. litorale* assembly is high for chelicerates, although more than twenty chromosomes are not unprecedented in arachnid taxa (Reyes-Lerma et al. 2021; You et al. 2024) and even higher numbers are known for some other aquatic arthropods, such as decapod crustaceans (P. Martin et al., 2016; Xu et al., 2021; Zhao et al., 2021; Zheng et al., 2024) (Suppl. Fig. 4A). While experimental confirmation via karyotyping is currently pending, the high BUSCO scores point towards a fairly complete genome, similar to other arthropods (Suppl. Fig. 4B). The size of the assembly lies in the previously reported range for chelicerates, albeit on the shorter side (Fig. 3A), with the only genomes that are consistently smaller being those of mites (Gregory & Young, 2020). The number of predicted protein-coding genes of *P. litorale* is modest compared to most other chelicerates (Fig. 3B), following the expected correlation with genome size (Fig. 3D).

Current chelicerate genome data suggest that they can be highly repetitive, with values routinely over 50-60% (Sanggaard et al., 2014; Sheffer et al., 2021) (Figs. 2, 3C). In this sense, the predicted repeat content of the *P. litorale* genome presented here (61.05%) is much more in line with what is already known from chelicerates than the *N. striatum* draft genome, which at just 7.14% is a clear outlier even in the context of all arthropods. However, while the *P. litorale* genome size and total repeat content are, on their own, within previously reported ranges for chelicerates and arthropods in general, the combination of high repeat content (>50%) in a small genome (<500Mb) is not (Fig. 3F). By demonstrating the frequent presence of repeat families in all raw reads we could exclude contamination as a likely source of the repetitive sequences, indicating that the remarkably high repeat/genome size ratio in *P. litorale* is real. Future studies will have to elucidate whether this represents a shared genomic feature of the entire pycnogonid lineage or rather a species-specific trait.

It is well-established that transposable elements (TEs) are potent drivers of genome evolution, offering a mutation source, providing raw material for novel cellular genes, enhancing rearrangements, duplicating or shuffling genes, and dispersing regulatory elements to new loci, among other functions (Bourque et al., 2018; Freeling et al., 2015). Moreover, there is a growing body of evidence that horizontal transfer of TEs between distantly related eukaryotes with close interactions (e.g., predator-prey, parasite-host) is more common than previously thought (Gilbert et al., 2010; Gilbert & Feschotte, 2018; Kambayashi et al., 2022). Given that *P. litorale* is an obligatory, tissue-sucking micropredator on selected sea anemone species, our cursory identification of similar repetitive motifs shared in the genomes of *P. litorale* and *M. metridium* thus warrants closer inspection and characterization of the unclassified repeats in the context of potential TE transfer.

Prior to our study, the microRNA families Mir-3931, Mir-5305, and Mir-5735 were classified as chelicerate-specific (Clarke et al., 2024). With data for pycnogonids lacking, however, only euchelicerate taxa could be included in previous microRNA surveys, leaving the question unresolved whether these three families evolved already at the base of the chelicerate tree. Interestingly, we found only evidence for Mir-3931s in the *P. litorale* genome. As the high score of >95% for conserved microRNAs speaks for a good representation of the total microRNA content in the assembly, this suggests that only Mir-3931 was already present in the last common ancestor of Chelicerata, whereas Mir-5305 and Mir-5735 are likely derived microRNAs of Euchelicerata.

### Novel genomic and transcriptomic resources unlock the *P. litorale* for chelicerate evo-devo

Molecular data for sea spiders have so far been scarce and taxonomically fragmented, hampering the interrogation of the pycnogonid body plan and its development with modern approaches. Until recently, gene expression data were limited to three Hox genes in *Endeis spinosa* larvae (Jager et al., 2006), while sequence resources were restricted to a draft genome for *Nymphon striatum* (Jeong et al., 2020) and to bulk transcriptomes for a handful of other species (Ballesteros et al., 2021). By sequencing the *P. litorale* genome and generating stage-specific transcriptomes (Suppl. Table 1) we provide here the most complete molecular description of pycnogonid development to date. This combines synergistically with pre-existing knowledge on laboratory husbandry (Brenneis et al., 2023; Vilpoux & Waloszek, 2003), ontogenesis and adult morphology (e.g. Alexeeva & Tamberg, 2021; Frankowski et al., 2022; Machner & Scholtz, 2010; Ungerer & Scholtz, 2009), as well as recently established hybridization chain reaction fluorescent in-situ hybridization (HCR-FISH) protocols in *P. litorale* (Klementz et al., 2024). Together, this resource expansion enables state-of-the-art studies on pycnogonids, paving the way to a better understanding of the evolution, development, and general biology of this ancient and phylogenetically important chelicerate lineage. Among others, the indirect developmental mode of pycnogonids promises new insight into the molecular underpinnings and evolution of chelicerate body patterning, which so far largely relies on a few arachnid species (e.g. Gainett et al., 2022; Oda & Akiyama-Oda, 2020) that exhibit direct development as one of the common, derived adaptations of terrestrialization. Beyond this, the high-quality genome of *P. litorale* represents an important milestone toward unlocking novel macrosynteny-based phylogenetic approaches (e.g. Schultz et al., 2023) for the interrogation of the recalcitrant higher-order relationships of Chelicerata.

### The Hox gene cluster of *P. litorale* provides evidence against a WGD in the chelicerate ancestor

The ancestral arthropod Hox cluster was likely comprised of ten genes (Pace et al., 2016), which contribute to governing segment identity along the anterior-posterior axis during development. So far, data on pycnogonid Hox genes have been fragmentary. The earliest, PCR-based survey of two species (*Nymphon gracile*, *Endeis spinosa*) did not retrieve a full Hox gene complement for either sea spider (Jager et al., 2006; Manuel et al., 2006), although, taking both species together, orthologs of all ten genes were present. While *abdA*/*Hox9* was not found in *E. spinosa*, only a highly divergent *abdA* sequence was identified in *N. gracile*. However, the draft genome of *N. striatum* (Jeong et al., 2020), a congener of *N. gracile*, shows no evidence of *abdA*, and neither do any of the hitherto generated developmental transcriptomes of other pycnogonid species (Ballesteros et al., 2021). In this study, we screened developmental transcriptomes of *P. litorale* and report its completely sequenced Hox cluster, the first for a sea spider (Fig. 4A). Given the lack of any *abdA* orthologs in our transcriptomic data and its absence from the intact Hox gene cluster, our results provide strong genomic evidence for a degradation/loss of *abdA* in the pycnogonid crown group. This inference receives additional support from the lack of IAB-4 in our microRNA survey. Together with Mir-10-P4 and Mir-10-P1, Iab-4 belongs to a conserved trio of microRNAs associated with the arthropod Hox cluster, in which Iab-4 is almost invariably located between *abdA* and *AbdB* (e.g. Pace et al., 2016; Yuan et al., 2024). Accordingly, the absence of a recognizable Iab-4 from the otherwise very complete suite of conserved microRNAs serves as additional indication for significant sequence degradation in the Hox cluster region that typically includes *abdA*. These novel findings in *P. litorale* call for a reinvestigation of the *N. gracile* Hox gene cluster with state-of-the-art sequencing approaches, as well as new genomic resources for Austrodecidae, the sister group of the remaining Pycnogonida, to test the inference that *abdA* loss is plesiomorphic for crown group sea spiders.

Given that Hox cluster duplications are commonly considered a strong indicator for WGD events (e.g. (Schwager et al., 2017; Sharma, 2023a)), the identification of a single, intact Hox cluster in *P. litorale* and the lack of additional Hox gene copies in any other part of the genome confidently anchors a non-duplicated genome in the chelicerate ground pattern (Fig. 4B). This unequivocally polarizes the non-duplicated genomes of apulmonate euchelicerate taxa as the ancestral condition and corroborates that the xiphosuran and arachnopulmonate WGDs are derived (Aase-Remedios et al., 2023; Sharma, 2023b).

### The reduced pycnogonid opisthosoma and the lack of *abdA* - cause or eHect?

The posterior Hox gene *abdA* is typically expressed in major parts of the chelicerate opisthosoma (Schwager et al., 2015; Turetzek et al., 2024) and it was previously hypothesized that the reduction of the pycnogonid opisthosoma may be linked to the loss or strong degeneration of this gene, leading to a loss of its patterning function (Manuel et al., 2006). Intriguingly, the same correlation is known for certain acariform mites with extremely reduced opisthosoma (Brückner et al., 2022; Grbić et al., 2011; Pace et al., 2016) (Fig. 4B), and, outside Chelicerata, in cirripede crustaceans, which likewise exhibit an extreme reduction of their posterior tagma, the abdomen (Deutsch & Mouchel-Vielh, 2003; Yuan et al., 2024). Losses of multiple trunk Hox genes, including *abdA*, are also associated with the compaction of the tardigrade body plan (Smith et al., 2016). Accordingly, the correlation of posterior tagma reduction and the absence of *abdA* across evolutionarily distant arthropod lineages may hint toward a second, higher-order function of *abdA* beyond specification of posterior segment identity, namely the formation and maintenance of posterior body segments. A precedent for such dual functions of Hox genes is known for *labial*/*Hox1* orthologs, which not only convey identity to the tritocerebral segment in insects and spiders, but are also crucial for its maintenance (Pechmann et al., 2015; Posnien & Bucher, 2010). If a similar dual function were to apply to *abdA*, its loss may have been sufficient to cause the reduction of posterior segments. In line with this scenario, an abrupt removal of *abdA* from the Hox cluster has been hypothesized for the body plan evolution of acariform mites (Pace et al., 2016). This was based on the reversed orientation of *AbdB*/*Hox10* compared to the rest of the genes in the Hox cluster, hinting at a potential chromosome inversion that impacted part of the posterior Hox genes.

Ideally, this proposed relationship would need to be tested via functional interrogation of *abdA*, something that has been hitherto unsuccessful in chelicerates (Sharma, 2023a). Outside of Chelicerata, however, functional insights were gained from several insects and one malacostracan crustacean, where *abdA* clearly instructs segment identity, whereas there is no evidence of its involvement in posterior segment formation and maintenance comparable to *lab* (Angelini et al., 2005; A. Martin et al., 2016; Stuart et al., 1993; Vachon et al., 1992). By extrapolation, this suggests that the loss of *abdA* function may also in chelicerates represent an insufficient condition for opisthosomal segment reduction. Moreover, unlike in acariform mites, the Hox cluster of *P. litorale* displays uniform orientation of all nine Hox genes present (incl. *AbdB*), disfavoring the idea of a chromosome inversion event that removed *abdA* from the cluster during sea spider evolution. Taken together, these observations suggest that the causal relationship may rather be reversed: the reduction of opisthosoma segments rendered their further specification obsolete, making a loss of *abdA* possible, for instance due to mutation accumulation to the point of extreme degradation and eventual pseudogenization. Notably, a similar scenario has been recently proposed to underlie the evolution of the compacted tardigrade body plan, which correlates with the absence of multiple Hox genes (Smith et al., 2024).

At the morphological level, fossils provide some support that the compaction of the segmented opisthosoma may have resembled a gradual process in the pycnogonid stem lineage (e.g. Sabroux et al., 2024; Siveter et al., 2023), suggestive of incremental instead of abrupt changes in the underlying developmental programs. However, it remains unknown to what extent crown group pycnogonids may have retained vestigial posterior segments during their development. While the transitory presence of up to two small posterior ganglia has been taken as evidence for this phenomenon (Brenneis et al., 2018; Brenneis & Scholtz, 2014), it awaits further confirmation by expression studies on the gene regulatory networks governing segmentation and segment identity in advanced postembryonic instars of *P. litorale*. Similar to corresponding studies on acariform mites and cirripedes (Barnett & Thomas, 2012, 2013; Blin et al., 2003; Gibert et al., 2000), such experiments will help clarifying whether the debated pycnogonid anal tubercle evolved by fusion of several vestigial opisthosomal segments, represents the terminal telson only, or is a composite structure of both elements (Brenneis et al., 2018, 2023; Brenneis & Scholtz, 2014; Dunlop & Lamsdell, 2017).

## Conclusions

In this study, we generated the first high-quality genome and comprehensive developmental transcriptomes for a sea spider, providing important reference points for studies into chelicerate evolutionary developmental biology and arthropod genome evolution. These novel resources will reinvigorate the systematic examination of pycnogonid development at the molecular level and afford the opportunity to polarize evolutionary developmental trajectories of body patterning at the base of the chelicerate tree. The single Hox gene cluster of *P. litorale* provides the first genomic evidence that no WGD events did occur already in the chelicerate stem lineage. Beyond this, it corroborates the lack of *abdA* in sea spiders, highlighting the independent evolutionary recurrence of a common genomic motif in distantly related arthropod groups that share a significant reduction of their posterior tagma.

## Author contributions

GB, AW, PPS conceived and planned the project. GB established and maintained the animal culture. GB collected and selected material for stage-specific RNAseq. CB, GB, EVWS, SSK, PPS performed protocol optimizations, dissections, and extractions for DNA and RNA sequencing. NP performed the genome assembly, scaffolding, genome annotation, orthology analysis. BF performed the microRNA analysis. PPS, SSK, EVWS provided PacBio and Omni-C data (Maine specimens) and additional developmental transcriptomes. NP, AW, GB wrote the first draft of the manuscript. All authors read, commented on, and approved the final version of the manuscript.

## Acknowledgements

The authors wish to thank Juan Montenegro for his help with genome assembly, contamination analysis, and genome annotation; Oleg Simakov for consulting about data generation and assembly methodology; Darrin Schultz for consulting about scaffolding and assembly fine-tuning; Marion Wanninger and Max Hämmerle for their support with the *P. litorale* husbandry. The computational results of this work have been achieved using the Life Science Compute Cluster (LiSC) of the University of Vienna.

## Supplemental Figures & Tables

**Suppl. Fig. 1:** - Comparison of COX1 (cytochrome oxidase I) sequences between lab culture (labelled “Helgoland”), the wildtype animals used (labelled “Maine”) and the NCBI entries MG934985, MG935177, MG935394, and HM425354 (*P. litorale* COX1 partial CDS). Genome alignment produced by Geneious v10.2.6.

**Suppl. Fig. 2:** Genome size estimation from k-mer (k=21) coverage statistics. A) GenomeScope profile for PacBio reads; B) GenomeScope profile for ONT reads; C) GenomeScope2 profile for PacBio reads; D) GenomeScope2 profile for ONT reads.

**Suppl. Fig. 3**: Visualization of the best hits for arachnid *abdA* sequences on pseudochromosome 56, in relation to the location of the *P. litorale Hox7/Antp* and *Hox8/Ubx* gene models. Notably, no hits are found between the gene models, where a presumptive *abdA* locus would be expected. For the accession IDs and sequences used for this, refer to Suppl. File 1, available on Zenodo (10.5281/zenodo.14185694).

**Suppl. Fig. 4:** Overview of A) (pseudo-)chromosome number and B) BUSCO completeness for different published arthropod genome assemblies. Each point represents one genome, with Hexapoda shown in yellow, chelicerates in magenta, myriapods in gray, and crustaceans in cyan. The bisected point shows the average of the distribution. The dashed red line denotes the values for *P. litorale* (this study). The underlying data can be found in Suppl. File 2 (10.5281/zenodo.14185694).

**Suppl. Table 1:** Overview of the sequencing data generated for the project. Technology: sequencing platform used. De novo: whether the data was used to assemble a de-novo transcriptome. Annotation: whether the transcriptomic data was used to predict protein-coding genes. Accession: European Nucleotide Archive Accession IDs.

**Suppl. Table 2:** Comparison of COX1 (cytochrome oxidase I) sequence identity between lab culture (labelled “Helgoland”), the wildtype animals used (labelled “Maine”) and the NCBI entries MG934985, MG935177, MG935394, and HM425354 (*P. litorale* COX1 partial CDS).

**Suppl. Table 3:** Iso-seq mixing strategy for the various developmental stages. We aimed for an approximately equimolar mix while trying to reach recommended concentrations for PacBio sequencing and considering the total amount of RNA extracted from each developmental stage. The RIN number is a score of RNA integrity (Schroeder 2006) with values ranging from 10 (intact) to 1 (totally degraded); DV200 denotes the percentage of RNA fragments with length greater than 200 nucleotides.

**Suppl. Table 4:** QUAST and BUSCO Arthropoda scores that document the progress of the Flye-based assembly.

**Suppl. Table 5:** Overview of the best hits for spider *abdA* sequences (also see Suppl. Fig. 3). The underlying sequences can be found in Suppl. File 1 (10.5281/zenodo.14185694).

**Suppl. Table 6:** List of NCBI BLAST hits for gene model r2_g3735, predicted to reside in the Hox cluster of *P. litorale*.

**Suppl. Table 7:** List of chelicerate reference genome assemblies found on NCBI that were at least scaffold level. Ticks and mites are overrepresented; when multiple species from the same genus were present, we chose the one with more genes, as generally most ticks and mites with sequenced genomes are parasitic and have reduced genomes. Among the remaining chelicerate taxa, spiders are overrepresented; here we chose the species with the least predicted gene models for each genus, in the hope that we would avoid false positives. We excluded *A. ventricosus* as it contains an uncharacteristic number of predicted proteins. The number of genes corresponds to the total number of genes in the annotation, not the protein-coding ones. Underlies Fig. 3.

**Suppl. Table 8:** Arthropod repeat content. Chelicerate, myriapod, and hexapod data as reported by Sheffer *et al*. in Table 4 (Sheffer 2021). Crustacean data as reported by Cui *et al*. (Cui 2021).

**Suppl. Table 9:** Chelicerate repeat content, broken down by common repeat families. Manually extracted from various publications.

**Suppl. Table 10:** presence/absence overview of miRNA families predicted in the *Pycnogonum litorale* genome by MirMachine and supplemented with manual curation.

